# Molecular control of cellulosic fin morphogenesis in ascidians

**DOI:** 10.1101/2023.09.08.556826

**Authors:** Maxence LANOIZELET, Christel ELKHOURY YOUHANNA, Sébastien DARRAS

## Abstract

The tunicates form a group of filter-feeding marine animals closely related to vertebrates. They share with them a number of features such as a notochord and a dorsal neural tube in the tadpole larvae of ascidians, one of the three groups that make tunicates. However, a number of typical chordate characters have been lost in different branches of tunicates, a diverse and fast-evolving phylum. Consequently, the tunic, a sort of exoskeleton made of extracellular material including cellulose secreted by the epidermis, is the unifying character defining the tunicate phylum. In the larva of ascidians, the tunic differentiates in the tail into a median fin (with dorsal and ventral extended blades) and a caudal fin. Here we have performed experiments in the ascidian *Phallusia mammillata* to address the molecular control of tunic 3D morphogenesis. We have demonstrated that the tail epidermis medio-lateral patterning essential for peripheral nervous system specification also controls tunic elongation into fins. More specifically, when tail epidermis midline identity was abolished by BMP signaling inhibition, or CRISPR/Cas9 inactivation of the transcription factor coding genes *Msx* or *Klf1/2/4/17*, median fin did not form. We postulated that this genetic program should regulate effectors of tunic secretion. We thus analyzed the expression and regulation in different ascidian species of two genes acquired by horizontal gene transfer (HGT) from bacteria, *CesA* coding for a cellulose synthase and *Gh6* coding for a cellulase. Although we have uncovered an unexpected dynamic history of these genes in tunicates, and high levels of variability in gene expression and regulation among ascidians, we showed that *Gh6* has a regionalized expression in the epidermis, and that it is a regulator of caudal fin formation. Our study constitutes an important step in the study of the integration of HGT-acquired genes into developmental networks, and a cellulose-based morphogenesis of extracellular material in animals.

## Introduction

The tunicates are filter-feeder sea animals, and are the closest invertebrate relatives of the vertebrates. They share with them a basic body plan most visible during embryonic life with the presence of a notochord and a dorsal hollow neural tube. Together with cephalochordates (amphioxus), vertebrates and tunicates form the chordates ^1,2^. A unique structure, the tunic, gives their name to the tunicates, unites them and distinguishes them from other chordates. It consists of a ’mantle’ of extracellular material secreted by the epidermis, that protects them from the outside world but also gives them their external appearance, shape and color. It can be best compared to the insect exoskeleton, the cuticle. Tunic formation is poorly known and studied. However, a major and striking discovery was the identification of cellulose as a major component of the tunic. This biomolecule is the most abundant on earth, and it is best known as the primary compound of plant cell wall with multiple applications in human life (paper, textiles, chemistry…). Tunicates are actually the only metazoans known to synthesize cellulose, a specificity that appeared through the acquisition of a *cellulose synthase* (*CesA*) gene from a bacteria through horizontal gene transfer (HGT) ^3–7^. This gene is expressed in the epidermis during embryogenesis, and is essential for cellulose production and proper metamorphosis. Importantly, in the absence of *CesA* a tunic still forms ^7^. This is in agreement with the fact that cellulose is not the sole component of the tunic that includes other extracellular material such as polysaccharides and proteoglycans ^8–12^. Very recently, another gene acquired through HGT, *Gh6*, has been described as expressed in the epidermis and possibly involved in cellulose metabolism and metamorphosis ^13,14^.

The tunic as mentioned above gives their shape to the tunicates. The most striking example comes from a the appendicularians (or larvaceans). Their tunic, also called the ’house’, has a sophisticated 3D structure with multiple parts and serves as a complex filtering and food - microalgae and bacteria - concentrating engine. Instead of cleaning the filter when clogged, appendicularians escape their house and build a new one, and they do so several times a day. This is an astonishing and fascinating case of 3D morphogenesis using extracellular matrix production that has been studied in some detail at the molecular level ^6,15–17^. We have chosen to turn to a much simpler case. Another group of tunicates, the ascidians (or sea squirts), possesses a tunic not only during the adult life but also during the larval life – a tadpole-like form. This larval tunic is longer along the dorsal, ventral and caudal parts of the tail to form dorsal, ventral and caudal fins. The median (dorsal and ventral) and caudal fins are thought to be essential for swimming ^18^. In the trunk, the tunic also differentiates with some specific elongations ^19^. We postulate that 3D morphogenesis of the larval ascidian tunic is regulated through specific secretion and/or modification of extracellular matrix components at the specific location where elongations/extensions occur. It is most likely that tunic shape is intimately linked to the patterning of the underlying epidermis. By studying the developmental mechanisms for the caudal peripheral nervous system (PNS) formation in the ascidian *Ciona intes=nalis*, we have uncovered cellular and molecular patterning of the tail epidermis into 3 main longitudinal rows of cells: the dorsal and ventral midlines, the four medio-lateral rows and the two lateral rows ^20^. Interestingly, the larval tunic shape is similar in distantly related species (Fig S1) ^21^, and the tail epidermis transcriptional patterning and the underlying gene regulatory network (GRN) is partly conserved ^22,23^.

Here, we have used the embryos of the ascidian *Phallusia mammillata* to interrogate the mechanisms of larval tunic morphogenesis. We showed that embryonic developmental mechanisms that pattern the tail epidermis also regulate tunic shape. Specifically, inhibition of BMP signaling that prevents ventral midline specification abolished ventral median fin formation. CRISPR/Cas9-mediated mutagenesis of *Msx* and *Klf1/2/4/17* coding for transcription factors expressed at the tail midlines impaired median fin elongation. We next sought for effectors of tunic elongation by focusing on HGT acquired cellulose-related genes, *CesA* and *Gh6*. We describe a dynamic evolutionary history of these genes in tunicates, and their expression patterns and regulation in four ascidian species using cross-species transcriptional assays. Despite a shared epidermal expression and a conserved regulation of *CesA* by the epidermis determinant Tfap2, the expression and transcriptional regulation of these HGT genes is unexpectedly variable. Finally, we uncovered a conserved regionalized epidermal expression of *Gh6* that is regulated by the tail epidermis GRN, and we propose that the cellulase encoded by *Gh6* restricts tunic elongation during fin blade formation.

## Material and methods

### Embryo obtention and manipulation

Adults from *Ciona intes=nalis* (formerly referred to *Ciona intes=nalis* type B ^24^) and *Ascidia mentula* were provided by the Centre de Ressources Biologiques Marines in Roscoff (EMBRC-France). *Phallusia mammillata* and *Molgula appendiculata* were provided by the Centre de Ressources Biologiques Marines in Banyuls-sur-mer (EMBRC-France). Gamete collection, embryo rearing and electroporation were performed as described previously ^23,25^. Staging of embryos is according to the developmental table of *Ciona robusta* ^26,27^. To improve tunic and fin blades formation amer dechorionation, embryos were cultured at 22°C (*P. mammillata*) or 19°C (*C. intes=nalis*) in the presence of 0.1% BSA on 1% agarose-coated dishes.

Dechorionated embryos were treated with 2.5 µM of the BMP receptor inhibitor DMH1 and with 150 ng/ml of recombinant BMP2 protein from early gastrula stages (St.10) as previously described ^22,28^. *In vivo* transcriptional assays were performed as described ^23,25^.

### Cellulosic larval fin visualization

When *P. mammillata* embryos reached the early swimming larval stage (St. 27; 16 hpf at 22°C), we transferred them in 2 ml low binding microtubes in 1 ml of sea water. They were fixed for 2 hrs at room temperature (RT) or overnight (O/N) at 4°C by adding 1 ml of fixation buffer (0.1 M MOPS, 0.5 M NaCl, 7.4% formaldehyde) to reach a final concentration of 3.7% of formaldehyde. Amer 3 successive washes in PBST (137 mM NaCl, 10 mM Na2HPO4, 2.68 mM KCl, 1.76 mM KH2PO4, 0.1% Tween20), we proceeded to the cellulose staining step.

Three different methods were optimized: (1) 2 hrs at RT or O/N at 4°C in 75 µg/ml of CBM3a-GFP (CZ00571, Nzytech), (2) 1 hr at RT in 0.1 % Calcofluor White (910090, Sigma-Aldrich), and (3) 3 hrs at RT in 0.1% Direct Red 23 (212490, Sigma-Aldrich), in PBST. Staining was stopped by 3 successive washes in PBST. Nuclei were stained with 0.5 μg/ml DAPI (62247, Fischer Scientific), or with 1 μM Sytox Green (S7020, Invitrogen) in PBST for 30 min at RT. CBM3a-GFP cellulose staining has been preferred since it did not produce non-specific staining of the larval body. Whole larvae were examined under a stereomicroscope (Olympus SZX16 equipped with a X-Cite Fluorescence Lamp Illuminators (Excelitas Technologies)), and imaged with a widefield inverted fluorescence microscope (Olympus model IX51HBO 100W equipped with a FL20 camera, Tucsen), and/or with a TCS SP8 X confocal microscope (Leica). 3D reconstruction and object modelling were conducted with the IMARIS microscopy imaging somware (Oxford Instrument).

### Molecular biology, gene identifiers and phylogeny

Gene identifiers, sequences and methods for RNA probe DNA templates and *cis*-regulatory DNA generation are described in Table S1. Genes and *cis*-regulatory regions were named according to the ascidian community nomenclature ^29^.

Putative Tfap2 binding sites were mapped on *CesA* loci using FIMO (hap://meme-suite.org/tools/fimo) ^30^ with matrices collected from the Jaspar database ^31^ and the GCCN3/4GGC motif ^32^. The sites were mutated by making the following changes G->C and C->G in order to keep the GC content unchanged (the validity of these changes was confirmed using FIMO). Wild type and mutated versions of the *Phmamm.CesA* promoter were synthesized as gBlocks (IDT).

Genomic regions that were amplified from genomic DNA using PCR or gBlocks were placed upstream of the *Ciinte*.*Fog* basal promoter and *LacZ* using the Gateway technology (Invitrogen) ^23,33^. Protein sequences for CesA and Gh6 were collected using blast from different genomic and transcriptomic resources listed in Table S2. All sequences were aligned using the MUSCLE program ^34^. Maximum-likelihood phylogenies were inferred using IQ-TREE (hap://iqtree.cibiv.univie.ac.at/) ^35^.

### In situ hybridization

Chromogenic *in situ* hybridization in the different ascidian species were performed using the protocol described earlier ^23,28^ with dig-labeled RNA probes described in Table S1.

### CRISPR/Cas9 gene inactivation

The procedure was adapted from a recent protocol based on microinjection of the RNP complex into *Phallusia* ^36^. Target sequences were designed with the help of CRISPOR (hap://crispor.tefor.net/) ^37^. We selected sequences with a high ’Doench score’ that target specific parts of the locus: one against the region coding for the N-terminal region of the protein and two against the region coding for functional domains. The sequences are shown in Table S3. *P. mammillata* unfertilized eggs were dechorionated ^25^ and injected with the following solution: 30 μM pgRNA (duplex between custom sequence-specific crRNA and universal tracrRNA (IDT), 18.6 μM Cas9 (1081058, IDT), 9 mM Hepes, 67.5 mM KCl, 2.2 mM CHAPS, 1.3% Fast Green, and 0.25% AlexaFluor555-dextran (D34679, Invitrogen) in nuclease free Duplex Buffer (IDT). For control embryos, a single pgRNA (against *Phmamm.Tyr* or *Brlanc.Ascl1/2.1* depending on the experiment) at 30 µM was used. For *Msx*, *Klf1/2/4/17* and *Gh6*, an equimolar mix of 3 pgRNAs totaling 30 µM was used. Injection was performed with a Femtojet micro-injector (Eppendorf) using needles pulled with a P-1000 Micropipette Puller (Sutter Instrument) starting with capillaries (0.1 OD x 0.78 ID x 100 L mm, GC100TF-10, Harvard Apparatus). One to three hours amer injection, eggs were fertilized and lem to develop at 22°C. Properly injected larvae were sorted using the AlexaFluor555-dextran fluorescence under a dissecting scope. Given the variability in tunic and fin formation from dechorionated embryos, a set of larvae injected with the control pgRNA was systematically generated and analyzed on the same batch of eggs as for the experimental pgRNAs targeting the gene of choice.

## Results

### Larval tunic formation in Phallusia mammillata

Previous studies on the molecular control of larval tunic formation have been performed on *Ciona robusta* (*Ciona intes=nalis* type A) ^4,7,13^. We thus first described larval tunic formation during *P. mammillata* development. Since tunic is transparent, we have used three dyes, Calcofluor White, DirectRed23 and CBM3-GFP (see Materials and Methods), to visualize the tunic. Although all dyes may not be specific to cellulose since they might bind other polysaccharides ^6^, they were useful to follow the course of tunic formation (Fig 1). Tunic was first seen at late tailbud stages (St. 24) as a thin layer surrounding the embryo (Fig 1A). It progressively thickened with a small caudal fin visible at St. 25 (Fig 1B). At hatching (St. 26), extended median and caudal fins were readily visible (Fig 1C,D).

**Figure 1.**
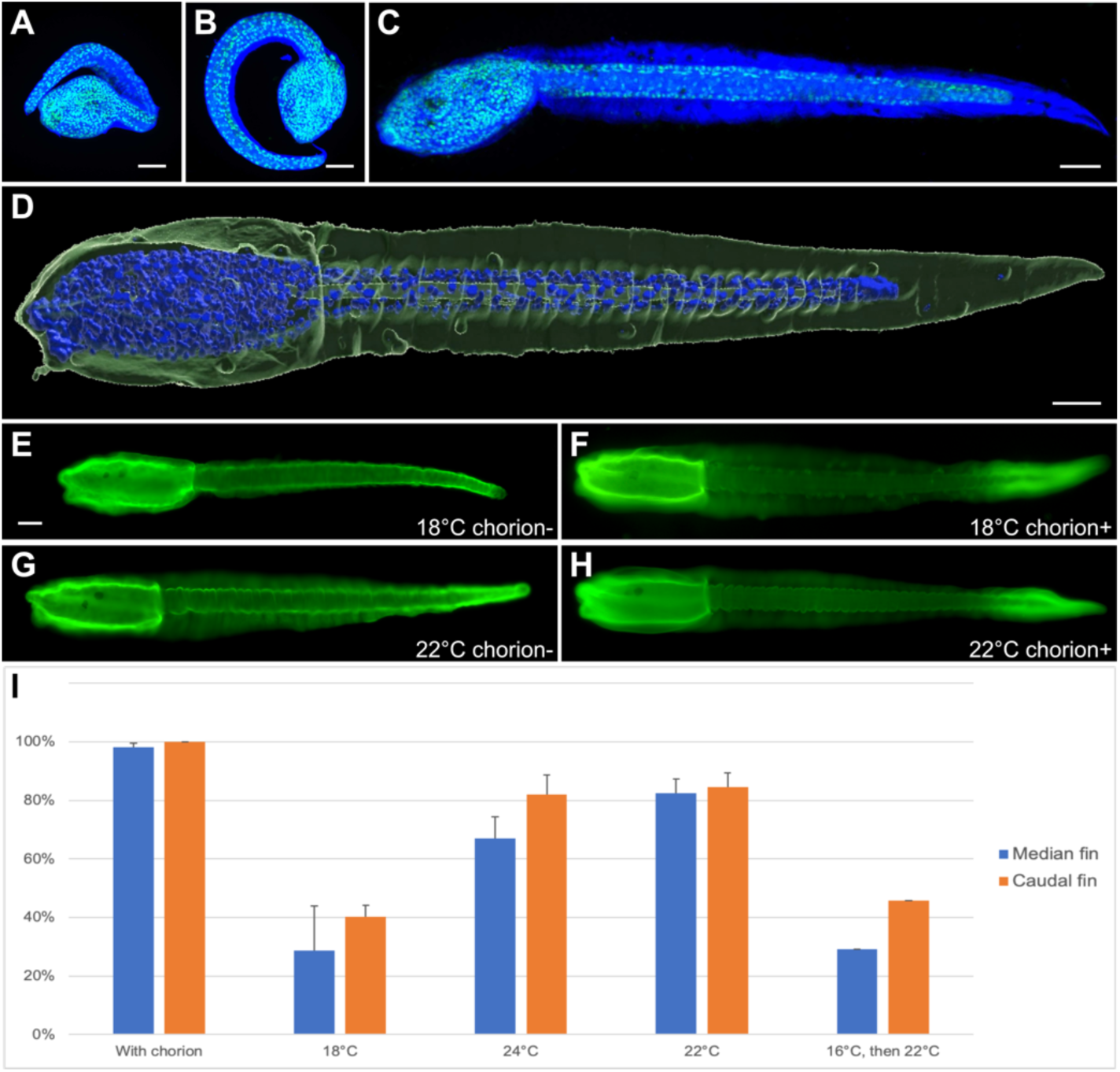
Tunic and fin blades development in *Phallusia mammillata*. **(A-C)** Embryos at St.24 (A) and St. 25 that developed in their chorions, and a hatching larva (C) were stained with Calcofluor White (blue) and Sytox Green (green). Maximum intensity projections from confocal z-stacks (the corresponding 3D visualizations can be found in Movies S1, S2 and S3). **(D)** 3D surface rendering from a confocal z-stack of a larva stained with CBM3-GFP (green) and DAPI (blue) (the corresponding 3D visualization can be found as Movie S4). **(E-H)** Effects of temperature during development on the formation of the tunic (stained with CBM3a-GFP in green) of larvae that arose from dechorionated eggs. Although the tunic of larvae that developed at 18°C was devoid of median and caudal fins most of the time (E), the tunic of larvae that developed at 22°C had more frequently median fins with a normal appearance and caudal fins with a reduced size (G). Chorionated embryos systematically gave rise to larvae with well-formed median and caudal fins (F,H). **(I)** Graph representing the proportions of larvae presenting median (blue) and caudal (orange) fins at the different culture temperatures (average values with error bars denoting standard deviation). With chorion: n=78 (3 experiments: one at 16°C, one at 18°C, and one at 24°C). 18°C without chorion: n=141 (3 independent experiments). 24°C without chorion: n=123 (3 independent experiments). 22°C without chorion: n=195 (4 independent experiments). 16°C, then 22°C without chorion: n=48 from a single experiment. Note that although the tunic was uniformly stained by CBM3a-GFP in dechorionated larvae, the tunic staining of chorionated larvae was always less intense in the middle part. Embryos in A and B have developed in their chorions, consequently they appear rolled up. Larvae in C-H are shown in lateral views with anterior to the left and dorsal to the top. Scale bars: 50 µm.

In order to manipulate ascidian embryos, the protective chorion is usually removed before or just amer fertilization. It has been known for long time that embryos deriving from such treatment are not fully normal. In particular lem/right asymmetry and tunic formation are disrupted ^21,38,39^. When we observed the tunic of larvae that derived from eggs whose chorion was removed before fertilization (using our routine protocol ^25^), the tunic was stained using cellulose dyes, but median and caudal fins were extremely reduced in size, abnormal or absent (Fig 1E,I) when compared with larvae that developed within their chorions (Fig 1F,I). We fortuitously found that increasing the temperature of embryonic development partly compensated for the absence of chorion. Dechorionated larvae that resulted from a development at 22°C or 24°C possessed fins more frequently (this frequency being quite variable from batch to batch), albeit usually reduced in size (Fig 1G,I). Interestingly, increasing the temperature before tunic and fin formation at early tailbud stages (St. 20/21) was not sufficient to recover fin formation (Fig 1I). This observation suggests that early events (possibly gene regulation), rather than a biophysical effect of temperature on tunic production, are the targets of temperature.

### Tail epidermis paIerning regulates median fin morphogenesis

Median fin blades elongate above a specific epidermal cell population, the tail neurogenic midlines (Fig 2A-C). We first inhibited the BMP signaling pathway that is necessary for ventral tail midline fate acquisition ^20,23,28^. Strikingly, treated larvae had no ventral fin (DMH1: 71% of larvae with median fin had only a dorsal fin, n=188; DMSO/BSA-control: 100% of larvae with median fin had both dorsal and ventral fins, n=181; results from four independent experiments; Fig 2D,E). A reciprocal experiment, treatment with recombinant BMP2 protein, leads to ectopic ventral midline fate in the entire tail epidermis ^22,23^. Contrary to our expectations, we did not observe a radial median fin all around the larval tail following such a treatment (Fig 2F). A median fin was not recognizable anymore and the tunic was somehow inflated making bulges all around the tail (this phenotype was observed in 99% of the embryos, n=272; results from three independent experiments). This observation is consistent with an excess of tunic production. We performed the same treatments on *C. intes=nalis* embryos (Fig S2). Although DMH1 treatment also led to a loss of ventral median fin, BMP2 treatment was different: the median fin was readily visible but numerous isolated fibers were seen protruding out of the tunic. While we are not yet able to explain the differences in phenotype for both species, the results argue for an excess of tunic production following BMP pathway activation and epidermis midline fate expansion.

**Figure 2.**
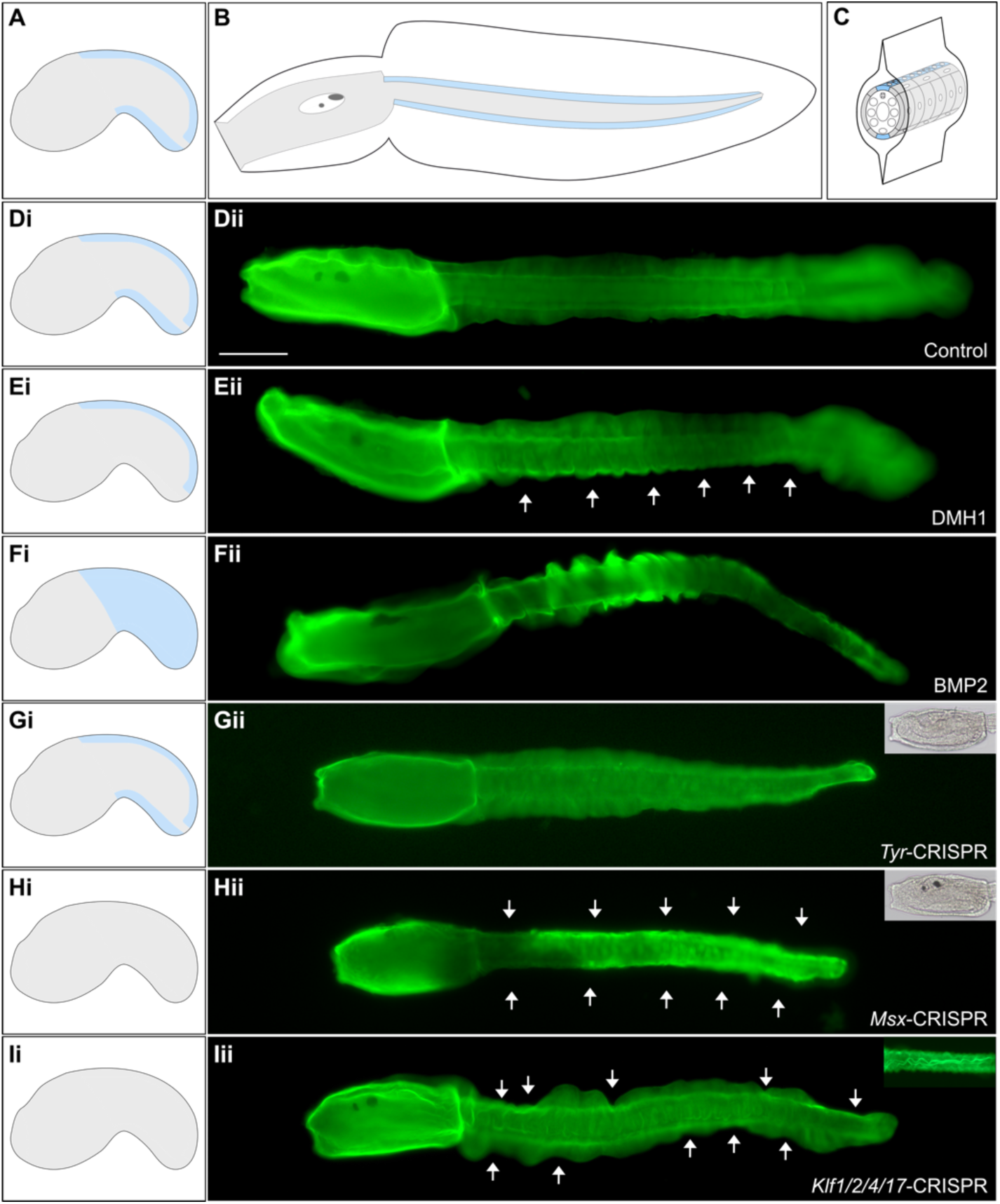
Molecular patterning of the tail epidermis regulates median fin formation. **(A-C)** Schematic representation of an early tailbud (A), a larva in lateral view (B) and a cross-section of the larval tail (C) with the embryo/larva itself in grey surrounded by the tunic. The tail epidermal neurogenic midlines that lie where median fins are positioned are highlighted in light blue. **(D-F)** Effects of BMP pathway modulation on median fin formation: control (BSA-and DMSO-treated larva) (Dii), DMH1-treated larva did not develop a ventral fin (Eii), and BMP2-treated larva with an excess of tunic making bulges (Fii). **(G-I)** Effects of CRISPR/Cas9 gene inactivation on tunic formation: *Tyr*-CRISPR larva formed a normal tunic with fins but lacked pigment cells (Gii), *Msx*-CRISPR larva lacked both ventral and dorsal median fins and had normal pigment cells (Hii), and *Klf1/2/4/17*-CRISPR larva presented indentations in the median fin that had an undulated shape (Iii). CBM3a-GFP staining of larvae in D-I (green). Insets in Gii and Hii: transmitted light picture of the trunk. Inset in Iii: dorsal view focused on the median fin. Larvae are shown in lateral view with anterior to the left and dorsal to the top. On the left panels (i): schematic representation of the distribution of the midline fate (light blue) in the different conditions. White arrows highlight the absence of median fin. Scale bar: 100 µm.

We next aimed at targeting downstream genes that belong to the tail PNS GRN using CRISPR/Cas9 gene editing. We used microinjection-mediated introduction of the ribonucleoprotein complex that has been recently described for *Phallusia* ^36^. As a control, we targeted *Tyrosinase* that is essential for melanin pigmentation of sensory organs found in the larval brain, the otolith and the ocellus. Absence of pigment cells was observed in 41% of larvae on average (59% with 2 pigment cells, 6% with 1 pigment cell, and 35% with no pigment cell; n=101 from three independent experiments; Fig 2G). We first examined the effects of mutating *Msx*, the gene coding for a homeodomain-containing transcription factor that lies at the top of the GRN ^22,40^. As anticipated, median fin formation was inhibited (*Tyr*-CRISPR: 79% of larvae with a median fin, n=78; *Msx*-CRISPR: 35% of larvae with a median fin, n=91; results from two independent experiments; Fig 2H). We then mutated *Klf1/2/4/17* that codes for a Zn-finger transcription factor acting downstream of *Msx* ^40^. Surprisingly, we did not phenocopy *Msx*-CRISPR larvae. The frequency of fin formation was similar to control *Tyr*-CRISPR larvae (*Tyr*-CRISPR: 87% median fin, 43% caudal fin, n=64; *Klf1/2/4/17*-CRISPR: 82% median fin, 46% caudal fin, n=52; results of two independent experiments). However, median fins were strongly malformed with a seemingly random local reduction in size leading to a wavy edge of the median fins (Fig 2I). This phenotype that was present in 77% of the larvae with median fins was never observed in *Tyr*-CRISPR, non-injected or non-dechorionated larvae. In addition, the median fin had an undulated shape, (Fig 2Iii inset). The results of this section demonstrate that acquisition of midline fate in the epidermis is essential for median fin formation.

### Phylogenetic distribution, embryonic expression and transcriptional regulation of cellulose-related HGT-acquired genes in ascidians

We hypothesized that elongation of the larval tunic into fin blades could be the result of specific regulation of cellulose production. We thus examined the phylogenetic distribution and embryonic expression of the two cellulose-related HGT-acquired genes *CesA* and *Gh6* in ascidians.

#### Pan epidermal expression of CesA

When searching for the ortholog of *CesA* in *P. mammillata* genome, we found two hits located on different scaffolds. Although predicted proteins differed in size, they had similar structures with 7 transmembrane domains, a glycosyl transferase 2 (GT2)/cellulose synthase domain and a glycosyl hydrolase 6 (GH6)/cellulase domain (Fig S3). Both genes had similar temporal dynamics (expressed during late embryonic development), but their levels of expression were highly different. To verify that the existence of these two genes was not an artifact of the *Phallusia* genome assembly, we surveyed tunicate genomic and transcriptomic data for *CesA* genes. As previously reported, we found a single *CesA* for most tunicates except *Oikopleura dioica* for which a lineage-specific duplication occurred ^6,41^. However, in addition to *P. mammillata*, we found three species that had 2 *CesA* genes (Fig S4, Table S2): *Phallusia fumigata*, *Ascidia mentula* and *Corella inflata*. Phylogenetic analysis demonstrated the existence of three clades (Fig S4): one that includes most previously described CesA, one with CesA proteins from *Oikopleura*, and one that includes the second copy present in these four species (we named these proteins CesA.b). The chromosomal assembly of *A. mentula* indicates that *CesA* and *CesA.b* are located on different chromosomes (Table S2), suggesting that recent local tandem duplication is not the source of *CesA.b* emergence.

We next determined the expression of *CesA* in three species: *P. mammillata*, *A. mentula* and *Molgula appendiculata* by *in situ* hybridization (Fig 3A-C). For all species, *CesA* had a pan-epidermal expression during embryonic development like in *Ciona* ^4,5,13^. However, in *M. appendiculata*, the expression was initiated much earlier (early gastrula stages) than in the other three species (late neurula stages). Despite *CesA.b* was expressed at much lower levels (Fig S3), we managed to detect its expression pattern during embryogenesis by *in situ* hybridization in *P. mammillata* (Fig 3D). *CesA.b* was broadly expressed in the epidermis of late tailbud embryos (St. 24) with weaker expression in the tail ventral and dorsal midlines.

**Figure 3.**
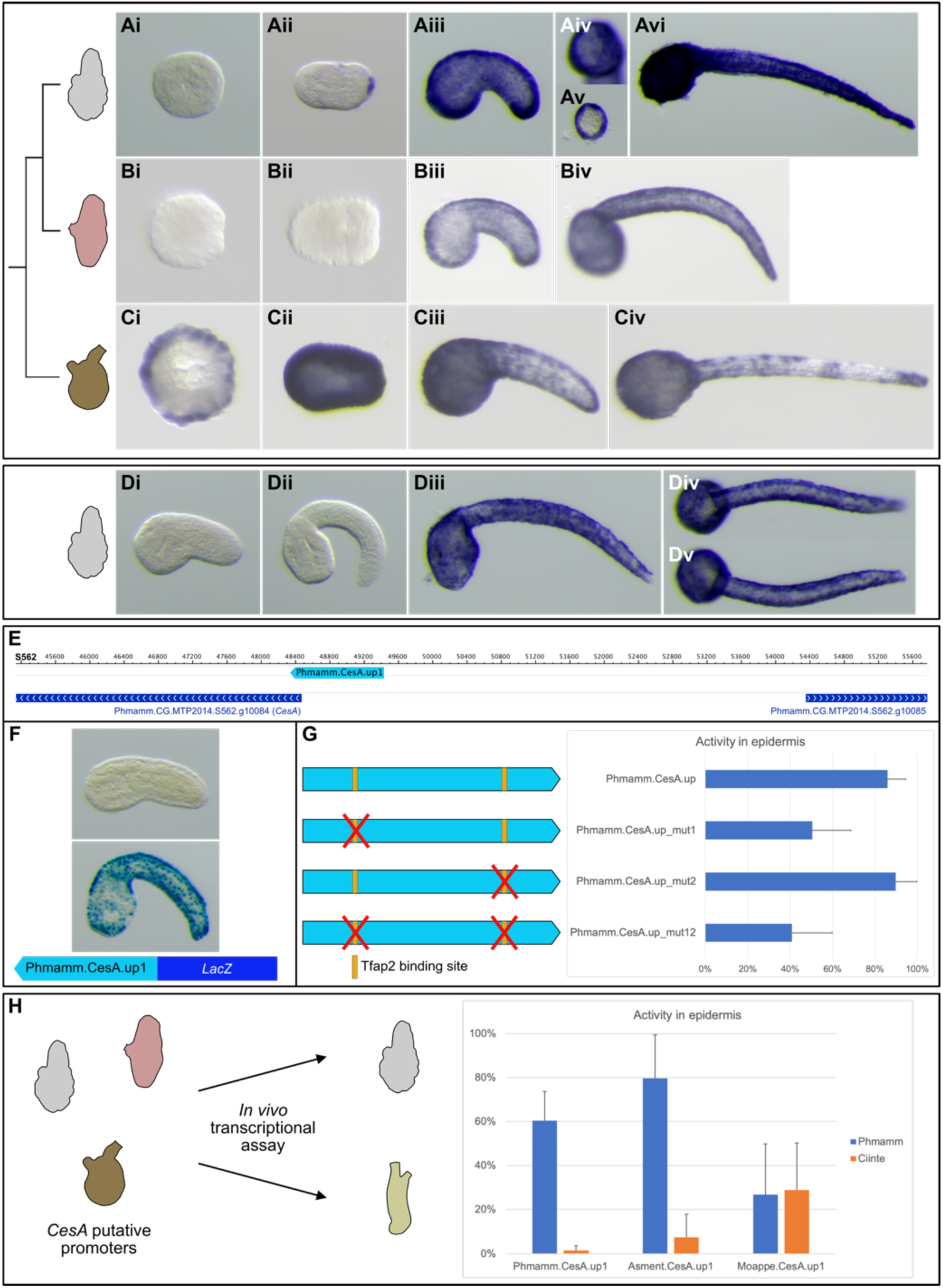
*CesA* expression and regulation in different ascidian species. **(A-C)** *In situ* hybridization for *CesA* during embryonic development of *Phallusia mammillata* (A), *Ascidia mentula* (B), and *Molgula appendiculata* (C). In *P.* mammillata, the expression was first detected in the caudal part of late neurulae (Aii). Expression was found throughout the epidermis with weaker staining in the palp forming region (Aiv). A cross-section of the tail shows that the staining was limited to the epidermis, and not found in internal tissues (Av). A similar pattern was found in *A. mentula* (B) and *M. appendiculata* (C) but with a much earlier onset of expression in the latter species (early gastrula: Ci). **(D)** *In situ* hybridization for *CesA.b* during embryonic development of *P. mammillata*. Transcripts were found throughout the epidermis from late tailbud stages (St. 24) with weaker expression in dorsal and ventral tail midlines (Diii-Dv). Embryos at the following stages are shown: gastrula (Ai,Bi,Ci), neurula (Aii,Bii,Cii), early/mid tailbud (Aiii-Av,Biii,Ciii,Di,Dii), and late tailbud (Avi,Biv,Civ,Diii-Dv). **(E-G)** Transcriptional regulation of *Phmamm.CesA* in the epidermis. A region of around 1 kb, Phmamm.CesA.up1, situated immediately upstream of *CesA* (E) was PCR-amplified and placed upstream of the *LacZ* reporter in inverted orientation. When tested in *P. mammillata* embryos (F), it was inactive at early tailbud stages (St. 19/20, no embryo stained in the epidermis, n=185, a single experiment) and active in the epidermis at late tailbud stages (St. 23/24, 60% of the embryos with epidermis staining, n=703 from 3 independent experiments). This region contains 2 putative Tfap2 binding sites (yellow bars in G). A synthetic version of Phmamm.CesA.up1, Phmamm.CesA.up, and 3 variants with Tfap2 mutations were placed in ’correct’ orientation upstream of *LacZ*, and their activity tested at mid-tailbud stages (St. 22) is summarized on the graph on the right in G (Phmamm.CesA.up, n=196; Phmamm.CesA.up_mut1, n=282; Phmamm.CesA.up_mut2, n=363; Phmamm.CesA.up_mut12, n=268; results from three independent experiments). **(H)** Swap assay of putative *CesA* promoters from three ascidian species. Each region was tested in both *P. mammillata* (Phmamm, blue) and *C. intestinalis* (Ciinte, orange), and the activity at mid/late tailbud stages in epidermis is displayed on the graph on the right (Phmamm.CesA.up1: in Phmamm n=703 from 3 experiments, in Ciinte n=348 from 2 experiments; Asment.CesA.up1: in Phmamm n=899 from 4 experiments, in Ciinte n=896 from 3 experiments; Moappe.CesA.up1: in Phmamm n=767 from 3 experiments, in Ciinte n=712 from 3 experiments). All pictures of embryos are lateral views with dorsal to the top and anterior to the left, except: animal views (Ai,Bi), vegetal view (Ci), neural plate views (Bii,Cii) with anterior to the left, frontal view (Aiv), dorsal view (Div), ventral view (Dv), and cross-section of the tail (Av).

#### Shared and divergent mechanisms regula=ng CesA expression in the epidermis

We isolated Phmamm.CesA.up1, a 1 kb genomic fragment immediately upstream of *Phmamm.CesA*, that prove sufficient to drive epidermal activity (Fig 3E,F). Since *Cirobu.CesA* is regulated by the transcription factor Tfap2-r.b ^42^, we searched for Tfap2 binding sites and identified two putative sites (Fig 3G). Mutagenesis followed by *in vivo* transcriptional assay indicated that the proximal site was dispensable (this site was actually predicted only by the GCCN3/4GGC motif, and not by the Jaspar matrices) and the distal site participated in *CesA* expression (Fig 3G). In fact, binding site mutation only diminished reporter activity whereas mutation of the single Tfap2 site fully abolished reporter activity in the *Ciona* experiment ^42^. To further probe potential differences between *Ciona* and *Phallusia*, we tested the transcriptional activity of Phmamm.CesA.up1 in *C. intes=nalis*. Surprisingly, it was inactive (Fig 3H). We isolated Asment.CesA.up1, a 1.6 kb long putative promoter from *A. mentula* that contains 3 predicted Tfap2 binding sites. Similarly to Phmamm.CesA.up1, this region was active in *Phallusia* epidermis but not in *Ciona* epidermis (Fig 3H). We next tested a candidate region from a distantly related species *M. appendiculata*. This 0.6 kb region containing 2 putative Tfap2 sites was equally active in both *Phallusia* and *Ciona* albeit at weaker levels than previous genomic fragments.

#### PaOerned epidermal expression of Gh6

We had identified the presence, in tunicate genomes, of another HGT gene originating from bacteria that codes for a most likely functional cellulase containing a transmembrane domain and an extracellular glycosyl hydrolase family 6 domain ^43^ that we named *Gh6*. Before the present study was completed, this finding was published by colleagues ^14^. Very recently, *Gh6* was described in *C. robusta* as being expressed in the epidermis during embryogenesis and as regulating larval tunic formation and metamorphosis ^13^. We confirmed the published phylogeny of tunicate Gh6 proteins (Fig S5). We identified a single genes for each species, except for the Thaliacean *Salpa thompsoni* that presented a lineage-specific duplication (Fig S5) ^14^.

We examined the expression pattern of *Gh6* during the embryonic development of 4 ascidian species. As expected, *Ciinte.Gh6* was expressed like *Cirobu.Gh6* (Fig 4A) ^13^ with first detectable expression in the epidermis of the Bp of the tail at late neurula/early tailbud stages (Fig 4Aii). Novel domains of expression in the epidermis appeared at late tailbud stages: palp region, dorsal epidermis at the trunk/tail junction and at the dorsal and ventral aspects of the tail (Fig 4Aiv). We found similar expression patterns in *P. mammillata* and *A. mentula* (Fig 4B,C). In *Phallusia*, we identified tail expressing cells as the 4 medio-lateral longitudinal rows of cells (2 ventral and 2 dorsal rows) ^20^ (Fig 4Biv-Bvi,Bviii). In the palp region, *Gh6* transcripts were depleted from future papillary protrusions (Fig 4Bvii). By contrast, *Moappe.Gh6* had a very different pattern (Fig 4D). Although it was expressed in the epidermis, the expression was detected earlier (neurula stages) and very broadly (entire epidermis with a depletion from the posterior tail as development proceeds).

**Figure 4.**
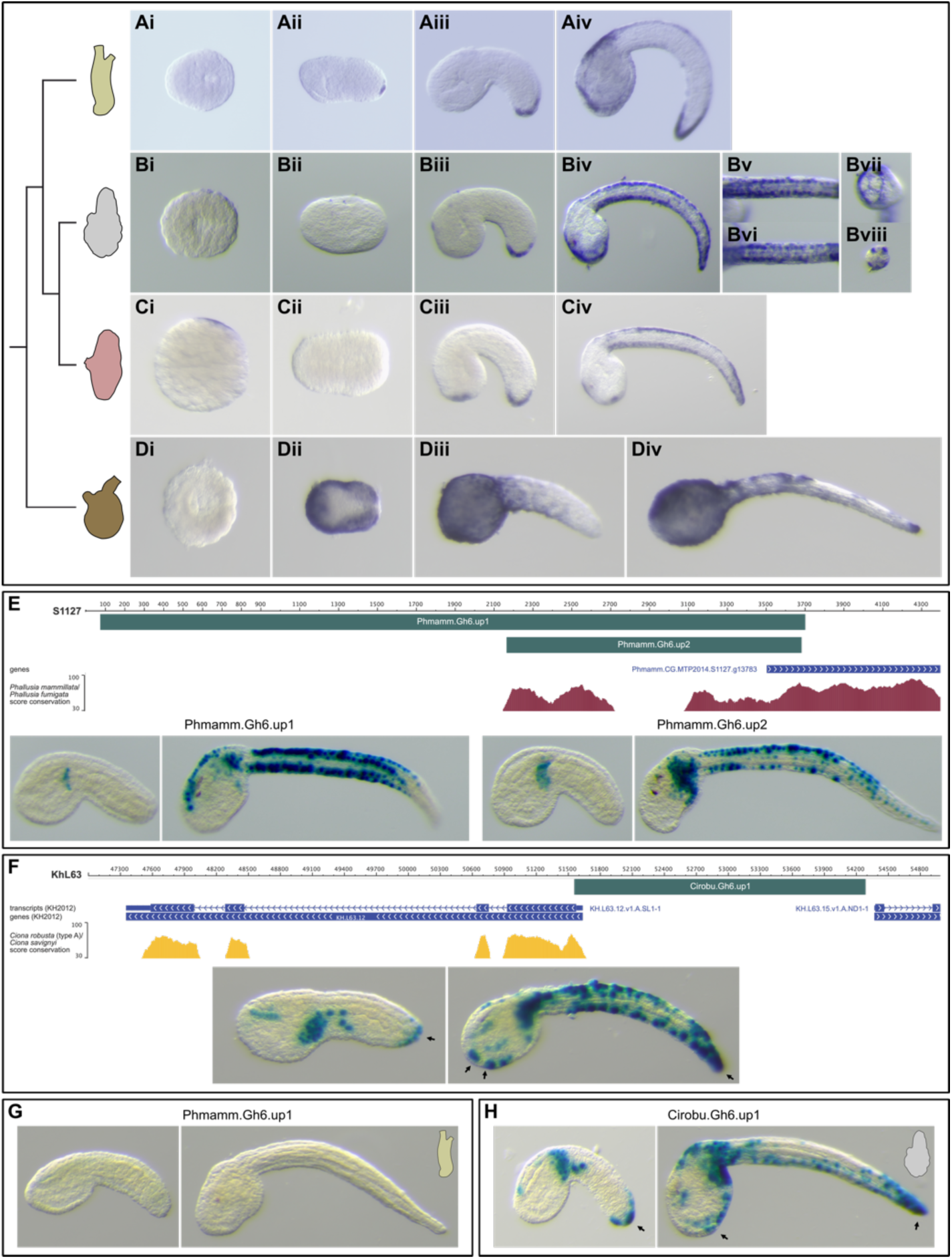
*Gh6* expression and regulation in different ascidian species. **(A-D)** *In situ* hybridization for *Gh6* during embryonic development of *Ciona intestinalis* (A), *Phallusia mammillata* (B), *Ascidia mentula* (C), and *Molgula appendiculata* (D). For the three phlebobranch species, *Gh6* was first detected at early tailbud stages in the epidermis at the tip of the tail. Then expression started anteriorly in the palp region and in the tail epidermis at late tailbud stages. Tail epidermis expression was detected in the four medio-lateral rows of cells (Bv,Bvi,Bviii). Expression in the palp region was likely absent from the future protruding papillae (Bvii). By contrast, *Moappe.Gh6* was expressed broadly in the epidermis starting at neurula stages (Dii-Div). Embryos at the following stages are shown: gastrula (Bi,Ci,Di), neurula (Ai,Bii,Cii,Dii), early/mid tailbud (Aii,Aiii,Biii,Ciii,Diii), and late tailbud (Aiv,Biv-Bviii,Civ,Div). **(E)** Transcriptional regulation of *Phmamm.Gh6* in the epidermis. A 3.6 kb region starting at the beginning of the scaffold 1127 immediately upstream of *Gh6*, Phmamm.Gh6.up1, was PCR-amplified, placed upstream of the *LacZ* reporter, and tested *in vivo*. It was active only in a part of the *Phmamm.Gh6* expression domain, the tail epidermal medio-lateral cells at late tailbud stages (St. 19-22: 3% of embryos with staining in this expression domain, n=401; St. 24-25: 75% of embryos with staining in this expression domain, n=521; results from four experiments). Phmamm.Gh6.up2, a 1.5 kb region embedded in Phmamm.Gh6.up1 containing conserved segments, had a similar activity (St. 19-22: 0% of embryos with this expression domain, n=670 from three experiments ; St. 24-25: 56% of embryos with this expression domain, n=556 from four experiments). **(F)** Transcriptional regulation of *Cirobu.Gh6* in the epidermis. Cirobu.Gh6.up1, a 2.7 kb region that almost corresponds to the entire upstream intergenic region, fully recapitulated *Gh6* expression with tail tip activity (arrows) detected in early tailbuds, and palp (arrows) and tail epidermis activity in late tailbuds (St. 19-22: 45% of embryos with expression in endogenous territories, n=300; St. 24-25: 73% of embryos with expression in endogenous territories, n=300; results from two experiments). **(G)** Phmamm.Gh6.up1 was not active in *C. intestinalis* embryos (n=367 from two experiments). **(H)** Cirobu.Gh6.up1 was active in *Gh6* expression domains in *P. mammillata* embryos (St. 21-22: 36% of embryos with expression in endogenous territories, n=448 from two experiments; St. 23-24: 72% of embryos with expression in endogenous territories, n=210 from three experiments). All pictures of embryos are lateral views with dorsal to the top and anterior to the left, except: animal views (Bi,Ci), vegetal view (Di), neural plate views (Ai,Cii,Dii) with anterior to the left, frontal view (Bvii), dorsal view (Bv), ventral view (Bvi), and cross-section of the tail (Bviii).

#### Shared and divergent mechanisms regula=ng Gh6 expression in the epidermis

To apprehend transcriptional regulation of *Gh6*, we isolated candidate *cis*-regulatory regions in both *Phallusia* and *Ciona*. In *Phallusia*, the largest region we could test (the *Gh6* locus lies at the edge of the currently available *P. mammillata* genome assembly MTP2014) reproduced late expression in the medio-lateral domains of the tail epidermis but not the palp and tail Bp regions (Fig 4E). A smaller fragment with DNA sequences conserved between *P. mammillata* and *P. fumigata* had a similar activity. In *Ciona*, the intergenic region upstream of *Gh6*, with no detectable DNA conservation between *C. robusta* and *C. savignyi*, recapitulated both the early tail Bp expression and the later palp and medio-lateral tail epidermis expression (Fig 4F). Surprisingly, although the regulatory elements isolated from *Phallusia* were inactive in *Ciona*, the elements from *Ciona* were active in *Phallusia* (Fig 4G,H). Altogether, the results from this section indicated an unexpected variability of *CesA* and *Gh6* at all level examined: phylogenetic distribution in tunicates, expression domains, and expression regulation. Nevertheless, these HGT-acquired genes are clearly deployed in the epidermis during late embryogenesis.

### Gh6 regulates caudal fin formation

As described previously (Fig 2), modifying tail epidermis patterning impacted fin blade formation. Since *CesA.b* and *Gh6* displayed a patterned expression in the epidermis, we determined their expression following BMP pathway modulation (Fig S6). Their expression patterns were modified in agreement with their site of expression: loss of ventral medio-lateral *Gh6* expression and ectopic *CesA.b* expression at the ventral midline when BMP was inhibited. We thus functionally evaluated the role of *Gh6* in tunic and fin blade formation. We selected this gene for two reasons. First, *Gh6* is expressed closely to where median and caudal fins emerge (Fig 4B). Second, by analogy with data from plants we postulated that *Gh6* may contribute to fin blade elongation through local increase of cellulose production ^44^. We thus inactivated *Gh6* using CRISPR/Cas9 (Fig 5). Contrary to our expectations, *Gh6* inactivation did not abolish fin elongation, but rather promoted an opposite fin phenotype. While the median fin was present similarly in both control-CRISPR larvae (in 73% of larvae, n=136 from two experiments) and in *Gh6*-CRISPR larvae (69%, n=155 from two experiments), the frequency of caudal fin presence increased from 48% in control-CRISPR larvae to 75% in *Gh6*-CRISPR larvae.

**Figure 5.**
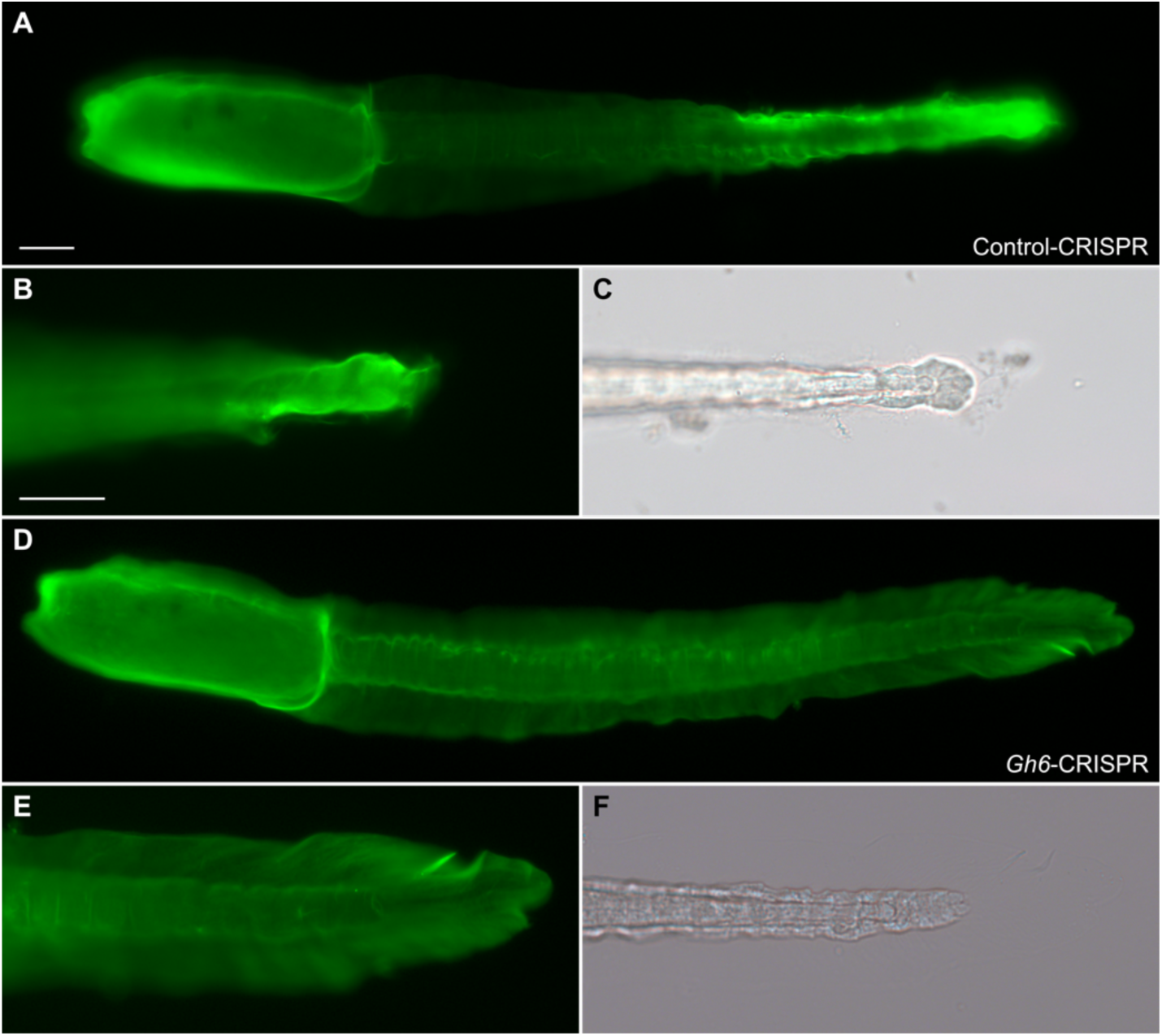
*Gh6* regulates caudal fin formation. Larvae resulting from microinjection of the CRISPR/Cas9 components targeting the *Brlanc.Ascl1/2.1* gene, a sequence absent from the *P. mammillata* genome (Control-CRISPR, A-C) or the *Phmamm.Gh6* gene (*Gh6*-CRISPR, D-F) were stained with CBM3a-GFP to visualize the tunic and fin blades. The caudal fin of the control larva ended where the tail ends (B,C). In contrast, the caudal fin of the *Gh6*-CRISPR larva was well developed. Scale bar: 50 µm.

## Discussion

### Function of Gh6 in tunic formation?

Cellulases have an obvious function of cellulose degradation during digestion for example. However, in both bacteria and plants, cellulases are part of the cellulose production machinery and promote the formation of crystalline over amorphous cellulose ^44,45^. For example, in *Arabidopsis*, KORRIGAN are cellulase mutants that present reduced biomass production. Although the enzymatic domains are different (GH8 in bacteria, GH9 in plants, and GH6 in tunicates), we postulated an analogous function for *Gh6* that would participate in local cellulose increased production and fin blade elongation. Instead of a reduction of fin blade elongation, we observed ’better’ caudal fin formation in our CRISPR experiment (Fig. 5). This suggest that *Gh6* normally prevents/regulates excessive fin elongation that would be triggered by another mechanism.

In a recent study, TALEN-mediated *Gh6* mutagenesis in *Ciona robusta* has also ascribed a function for *Gh6* in tunic formation with an excess of tunic where sensory papillae normally protrude ^13^. In addition, the formation of these papillae was prevented. This is an interesting phenotype given the regionalized expression of *Gh6* at this location (Fig 4). In our experiments, we did not observe any phenotype in the palps. The difference could stem from species-specific functions of *Gh6*, but the specificity of the reported phenotype in *Ciona* is questioned by the authors themselves since it could stem from toxicity of the TALEN system ^13^. It is also important to note that fin formation has not been examined in the cited study.

We were surprised by the modest phenotype for *Gh6*-CRISPR. A hint at the non-essential role of *Gh6* in fin formation comes from our description of its expression in different species (Fig 4). In *Molgula*, *Gh6* was found epidermal but without regionalized expression. It is thus unlikely to regulate local tunic expansion into fin blades. While this observation could stem from an inherent divergence in this species which belongs to a very fast-evolving genus, a better candidate would be a gene with conserved regionalized expression in all ascidians that make fin blades (Fig S1). Tunicate genomes encode a number of glycosyl hydrolases (annotated for *Phallusia* in the CAZY database ^43^), and one of them might fulfill the requirements for our above model for fin elongation. We are currently searching for genes coding for cellulase expressed in the epidermis during late embryogenesis in different ascidian species.

### Cellulose-related HGT-acquired genes in ascidians: dynamic evolutionary history and developmental system driR

HGT-based capacity of producing cellulose is at the core of tunic formation, a defining feature of the tunicate phylum. As discussed in previous reports ^4,6,14,41,42^, the process is not limited to the initial ancient cross-kingdom gene transfer. Beyond the open question of the number of HGT events (Have *CesA* and *Gh6* been acquired simultaneously? What about other yet to be described HGT-acquired genes?), such events imply novel functionalities for the host, hence maintenance of the gene and control of its expression. We discuss the integration of *CesA* and *Gh6* into developmental GRNs in the next section. Here, we would like to emphasize the surprisingly elevated degree of variations we have uncovered at different levels: evolutionary history, gene expression, and gene regulation, despite very similar tunic and fins in ascidian tadpole larvae. We have discovered a novel lineage-specific duplication of *CesA* in the ascidian sub-lineage comprising *Phallusia* and *Ascidia*. A similar case has been described in the appendicularian *Oikopleura dioica* where the two paralogs are deployed during separate parts of the life cycle and in different body parts (*CesA1* during the embryonic phase in the lateral tail epidermis and *CesA2* during the adult phase in the trunk epidermis). Since we have no data about gene expression outside the embryonic period in *Phallusia*, we are not able to tell whether we are in presence of a similar scenario. We could nevertheless describe co-expression of *CesA* and its paralog *CesA.b* in the epidermis of late embryos. The functional relevance of this observation and the regionalized expression of *CesA.b* will await further experiments. We have observed similar expression patterns for *CesA* and *Gh6* expression in the three phlebobranch species, but striking differences for *Molgula*, a stolidobranch ascidian (Fig 3,4). This could be, as discussed above and earlier ^23^, that *Molgula* is an outlier. This will require further documentation on gene expression at broader scale in ascidians. Nevertheless, this is very suggestive that even phylum-defining components are subject to drim and are very plastic. More surprisingly, we uncovered differences in transcriptional regulation for both genes between *Ciona* and *Phallusia*: promoters of *Phallusia* were not active in *Ciona*, whereas the *Gh6* promoter of *Ciona* was active in *Phallusia*. This *’asymmetric intelligibility’* is a situation that we have not observed when we analyzed a number of developmental regulators ^23^, but that has been reported between *Ciona* and *Molgula*, species that have separated a longer time ago ^46^.

### Integration of the tunic production machinery to the developmental genetic programs

By studying tail PNS formation, we had uncovered medio-lateral cellular and molecular epidermis patterning ^20^. Here, we show that the early inducing cue BMP signaling, and downstream nodes of the GRN, *Msx* and *Klf1/2/4/17*, also regulate median fin formation (Fig 2). It would thus be important to understand how HGT-acquired genes or yet unidentified effectors of fin elongation are integrated into the tail PNS GRN by dissecting the *cis*-regulatory regions of genes such as *Gh6*. Importantly, the tail PNS GRN that we and others have been deciphering ^20,22,23,40,47–50^ most likely diverges at some point into a ’neuronal GRN’ and a ’fin blade GRN’ since the tail midlines give rise to both peripheral sensory neurons and fins. Comparative work among ascidians should help pointing out these sub-networks. We have identified divergent expression patterns for nodes of the PNS GRN ^22,23^: some genes were expressed throughout the midlines in all species examined whereas some genes were expressed only in a sub-domain in some species. It turned out that, in these latter species, peripheral neurons do not form all along the midlines but only in parts of them. This is thus most likely that ’pan-midline’ genes fit into the ’fin blade GRN’ (all ascidians larvae have a median fin), and that genes with divergent expression fit into the ’neuronal GRN’. This will be an exciting line of comparative functional exploration.

### Cellulose, extracellular material and fin blade 3D morphogenesis

The molecular control of tunic and fin formation is yet underexplored. There are a number of open questions arising from previous studies and the present manuscript. Cellulose is an essential emblematic component of the tunic. However, biochemical studies indicate that other polysaccharides and extracellular matrix components, in particular glycoproteins, make up the tunic. Moreover, *CesA* inactivation clearly shows that without cellulose a tunic still forms but with an abnormal fin ^7^. In addition, *CesA* alone is unlikely sufficient for cellulose synthesis since a number of molecular partners are required ^4^.

The larval cellulosic fins are essential for swimming. Consequently, they are likely essential for a key life cycle transition: the larval dispersal and most importantly the settlement before metamorphosis and the switch to a sessile form. Re-investigating tunic composition through the angle of molecular EvoDevo sounds a promising approach to understand extracellular tunic 3D morphogenesis into fin blades by combining developmental GRN data with scRNA-seq data and CRISPR/Cas9 gene inactivation studies in several ascidian species with sequenced and annotated genomes.

## Conflict of interest

The authors declare that they have no conflict of interest.

## Acknowledgements

We thank G. Diaz (Port-Vendres) and staff (M. Fuentes, divers and boat crew) at the marine stations of Banyuls-sur-Mer and Roscoff (French node of the European research infrastructure EMBRC) for providing animals. We would like to acknowledge the BioPiC imaging facility (Sorbonne Université/CNRS, Banyuls-sur-mer) for access to microscopes. We acknowledge H. Yasuo for the gim of cDNA clones, A. McDougall for sharing the CRISPR protocol before publication and R. Chowdhury for establishing and teaching the ’Femtojet *Phallusia* injection’ procedure in our team.

## Funding

SD is CNRS staff. This work was supported by CNRS and Sorbonne Université, and by specific grants from the CNRS (DBM2020 and DBM2022 from INSB) and from Sorbonne Université (*AnimalCellulose* project within the Emergence 2021 framework).

## Authors’ contributions

ML and SD designed the project and wrote the manuscript. ML, CEY and SD performed experiments. SD obtained funding and supervised the project. All authors edited the manuscript, read and approved the final version.

## Data availability

All data generated or analyzed during this study are included in the manuscript and supporting files.

## Notes

### Competing Interest Statement

The authors have declared no competing interest.

## References

1. Satoh, N. (1994). Developmental biology of ascidians (Cambridge University Press).

2. Lemaire, P. (2011). Evolutionary crossroads in developmental biology: the tunicates. Development 138, 2143–2152. 10.1242/dev.048975.

3. Dehal, P., Satou, Y., Campbell, R.K., Chapman, J., Degnan, B., De Tomaso, A., Davidson, B., Di Gregorio, A., Gelpke, M., Goodstein, D.M., et al. (2002). The dram genome of Ciona intestinalis: insights into chordate and vertebrate origins. Science 298, 2157–2167.

4. Maahysse, A.G., Deschet, K., Williams, M., Marry, M., White, A.R., and Smith, W.C. (2004). A functional cellulose synthase from ascidian epidermis. Proc Natl Acad Sci U S A 101, 986–991.

5. Nakashima, K., Yamada, L., Satou, Y., Azuma, J., and Satoh, N. (2004). The evolutionary origin of animal cellulose synthase. Dev Genes Evol 214, 81–88.

6. Sagane, Y., Zech, K., Bouquet, J.M., Schmid, M., Bal, U., and Thompson, E.M. (2010). Functional specialization of cellulose synthase genes of prokaryotic origin in chordate larvaceans. Development 137, 1483–1492. 10.1242/dev.044503.

7. Sasakura, Y., Nakashima, K., Awazu, S., Matsuoka, T., Nakayama, A., Azuma, J., and Satoh, N. (2005). Transposon-mediated insertional mutagenesis revealed the functions of animal cellulose synthase in the ascidian Ciona intestinalis. Proc Natl Acad Sci U S A 102, 15134–15139.

8. Lübbering-Sommer, B., Compère, P., and Goffinet, G. (1996). Cytochemical investigations on tunic morphogenesis in the sea peach Halocynthia papillosa (Tunicata, Ascidiacea). 1: demonstration of polysaccharides. Tissue Cell 28, 621–630.

9. Lübbering-Sommer, B., Compère, P., and Goffinet, G. (1996). Cytochemical investigations on tunic morphogenesis in the sea peach Halocynthia papillosa (Tunicata, Ascidiacea) 2: demonstration of proteins. Tissue and Cell 28, 651–661. 10.1016/S0040-8166(96)80069-3.

10. Pavao, M.S., Aiello, K.R., Werneck, C.C., Silva, L.C., Valente, A.P., Mulloy, B., Colwell, N.S., Tollefsen, D.M., and Mourao, P.A. (1998). Highly sulfated dermatan sulfates from Ascidians. Structure versus anticoagulant activity of these glycosaminoglycans. J Biol Chem 273, 27848–27857.

11. Smith, M.J., and Dehnel, P.A. (1971). The composition of tunic from four species of ascidians. Comparative Biochemistry and Physiology Part B: Comparative Biochemistry 40, 615–622. 10.1016/0305-0491(71)90136-2.

12. Smith, M.J., and Dehnel, P.A. (1970). The chemical and enzymatic analyses of the tunic of the ascidian Halocynthia aurantium (pallas). Comparative Biochemistry and Physiology 35, 17–30. 10.1016/0010-406X(70)90909-6.

13. Li, K.-L., Nakashima, K., Hisata, K., and Satoh, N. (2023). Expression and possible functions of a horizontally transferred glycosyl hydrolase gene, GH6–1, in Ciona embryogenesis. EvoDevo 14, 1–14. 10.1186/s13227-023-00215-x.

14. Li, K.-L., Nakashima, K., Inoue, J., and Satoh, N. (2020). Phylogenetic Analyses of Glycosyl Hydrolase Family 6 Genes in Tunicates: Possible Horizontal Transfer. Genes 11, 937. 10.3390/genes11080937.

15. Kishi, K., Hayashi, M., Onuma, T.A., and Nishida, H. (2017). Patterning and morphogenesis of the intricate but stereotyped oikoplastic epidermis of the appendicularian, Oikopleura dioica. Developmental Biology 428, 245–257. 10.1016/j.ydbio.2017.06.008.

16. Sagane, Y., Hosp, J., Zech, K., and Thompson, E.M. (2010). Cytoskeleton-mediated templating of complex cellulose-scaffolded extracellular structure and its association with oikosins in the urochordate Oikopleura. Cell Mol Life Sci. 10.1007/s00018-010-0547-8.

17. Ganot, P., and Thompson, E.M. (2002). Patterning through differential endoreduplication in epithelial organogenesis of the chordate, Oikopleura dioica. Developmental biology 252, 59–71.

18. McHenry, M.J. (2005). The morphology, behavior, and biomechanics of swimming in ascidian larvae. Can. J. Zool. 83, 62–74.

19. Sato, Y., Terakado, K., and Morisawa, M. (1997). Test cell migration and tunic formation during posthatching development of the larva of the ascidian, Ciona intestinalis. Development, Growth & Differentiation 39, 117–126. 10.1046/j.1440-169X.1997.00013.x.

20. Pasini, A., Amiel, A., Rothbacher, U., Roure, A., Lemaire, P., and Darras, S. (2006). Formation of the Ascidian Epidermal Sensory Neurons: Insights into the Origin of the Chordate Peripheral Nervous System. PLoS Biol 4, e225.

21. Cloney, R.A. (1990). Larval Tunic and the Function of the Test Cells in Ascidians. Acta Zoologica 71, 151–159. 10.1111/j.1463-6395.1990.tb01190.x.

22. Chowdhury, R., Roure, A., le Pétillon, Y., Mayeur, H., Daric, V., and Darras, S. (2022). Highly distinct genetic programs for peripheral nervous system formation in chordates. BMC Biol 20, 1–25. 10.1186/s12915-022-01355-7.

23. Coulcher, J.F., Roure, A., Chowdhury, R., Robert, M., Lescat, L., Bouin, A., Carvajal Cadavid, J., Nishida, H., and Darras, S. (2020). Conservation of peripheral nervous system formation mechanisms in divergent ascidian embryos. eLife 9, e59157. 10.7554/eLife.59157.

24. Brune[, R., Gissi, C., Pennati, R., Caicci, F., Gasparini, F., and Manni, L. (2015). Morphological evidence that the molecularly determined Ciona intes=nalis type A and type B are different species: *Ciona robusta* and *Ciona intes=nalis*. Journal of Zoological Systematics and Evolutionary Research 53, 186–193. 10.1111/jzs.12101.

25. Darras, S. (2021). En masse DNA Electroporation for in vivo Transcriptional Assay in Ascidian Embryos. Bio-protocol 11, e4160.

26. Hotta, K., Mitsuhara, K., Takahashi, H., Inaba, K., Oka, K., Gojobori, T., and Ikeo, K. (2007). A web-based interactive developmental table for the ascidian *Ciona intes=nalis*, including 3D real-image embryo reconstructions: I. From fertilized egg to hatching larva. Developmental Dynamics 236, 1790–1805. 10.1002/dvdy.21188.

27. Hotta, K., Dauga, D., and Manni, L. (2020). The ontology of the anatomy and development of the solitary ascidian Ciona : the swimming larva and its metamorphosis. Scientific Reports 10, 17916. 10.1038/s41598-020-73544-9.

28. Roure, A., Chowdhury, R., and Darras, S. (2023). Regulation of anterior neurectoderm specification and differentiation by BMP signaling in ascidians. Development 150, dev201575. 10.1242/dev.201575.

29. Stolfi, A., Sasakura, Y., Chalopin, D., Satou, Y., Christiaen, L., Dantec, C., Endo, T., Naville, M., Nishida, H., Swalla, B.J., et al. (2015). Guidelines for the nomenclature of genetic elements in tunicate genomes. genesis 53, 1–14. 10.1002/dvg.22822.

30. Grant, C.E., Bailey, T.L., and Noble, W.S. (2011). FIMO: scanning for occurrences of a given motif. Bioinformatics 27, 1017–1018. 10.1093/bioinformatics/btr064.

31. Fornes, O., Castro-Mondragon, J.A., Khan, A., van der Lee, R., Zhang, X., Richmond, P.A., Modi, B.P., Correard, S., Gheorghe, M., Baranašić, D., et al. (2020). JASPAR 2020: update of the open-access database of transcription factor binding profiles. Nucleic Acids Res 48, D87–D92. 10.1093/nar/gkz1001.

32. Eckert, D., Buhl, S., Weber, S., Jäger, R., and Schorle, H. (2005). The AP-2 family of transcription factors. Genome Biol 6, 1–8. 10.1186/gb-2005-6-13-246.

33. Roure, A., Rothbacher, U., Robin, F., Kalmar, E., Ferone, G., Lamy, C., Missero, C., Mueller, F., and Lemaire, P. (2007). A mulBcasseae Gateway vector set for high throughput and comparative analyses in ciona and vertebrate embryos. PLoS ONE 2, e916.

34. Edgar, R.C. (2004). MUSCLE: multiple sequence alignment with high accuracy and high throughput. Nucleic Acids Research 32, 1792–1797. 10.1093/nar/gkh340.

35. Trifinopoulos, J., Nguyen, L.-T., von Haeseler, A., and Minh, B.Q. (2016). W-IQ-TREE: a fast online phylogenetic tool for maximum likelihood analysis. Nucleic Acids Research 44, W232–W235. 10.1093/nar/gkw256.

36. McDougall, A., Hebras, C., Gomes, I., and Dumollard, R. (2021). Gene Editing in the Ascidian Phallusia mammillata and Tail Nerve Cord Formation. In Developmental Biology of the Sea Urchin and Other Marine Invertebrates: Methods and Protocols Methods in Molecular Biology., D. J. Carroll and S. A. Stricker, eds. (Springer US), pp. 217–230. 10.1007/978-1-0716-0974-3_13.

37. Concordet, J.-P., and Haeussler, M. (2018). CRISPOR: intuitive guide selection for CRISPR/Cas9 genome editing experiments and screens. Nucleic Acids Research 46, W242– W245. 10.1093/nar/gky354.

38. Kourakis, M.J., Bostwick, M., Zabriskie, A., and Smith, W.C. (2021). Disruption of lem-right axis specification in Ciona induces molecular, cellular, and functional defects in asymmetric brain structures. BMC Biology 19, 141. 10.1186/s12915-021-01075-4.

39. Shimeld, S.M., and Levin, M. (2006). Evidence for the regulation of lem-right asymmetry in Ciona intestinalis by ion flux. Dev Dyn 235, 1543–1553.

40. Roure, A., and Darras, S. (2016). Msxb is a core component of the genetic circuitry specifying the dorsal and ventral neurogenic midlines in the ascidian embryo. Developmental Biology 409, 277–287. 10.1016/j.ydbio.2015.11.009.

41. Inoue, Nakashima, and Satoh (2019). ORTHOSCOPE Analysis Reveals the Presence of the Cellulose Synthase Gene in All Tunicate Genomes but Not in Other Animal Genomes. Genes 10, 294. 10.3390/genes10040294.

42. Sasakura, Y., Ogura, Y., Treen, N., Yokomori, R., Park, S.-J., Nakai, K., Saiga, H., Sakuma, T., Yamamoto, T., Fujiwara, S., et al. (2016). Transcriptional regulation of a horizontally transferred gene from bacterium to chordate. Proceedings of the Royal Society B: Biological Sciences 283, 20161712. 10.1098/rspb.2016.1712.

43. Drula, E., Garron, M.-L., Dogan, S., Lombard, V., Henrissat, B., and Terrapon, N. (2022). The carbohydrate-active enzyme database: functions and literature. Nucleic Acids Research 50, D571–D577. 10.1093/nar/gkab1045.

44. Cosgrove, D.J. (2005). Growth of the plant cell wall. Nature reviews 6, 850–861.

45. Noack, L.C., and Persson, S. (2023). Cellulose synthesis across kingdoms. Current Biology 33, R251–R254. 10.1016/j.cub.2023.01.044.

46. Stolfi, A., Lowe, E.K., Racioppi, C., Ristoratore, F., Brown, C.T., Swalla, B.J., and Christiaen, L. (2014). Divergent mechanisms regulate conserved cardiopharyngeal development and gene expression in distantly related ascidians. eLife 3, e03728. 10.7554/eLife.03728.

47. Imai, K.S., Levine, M., Satoh, N., and Satou, Y. (2006). Regulatory blueprint for a chordate embryo. Science (New York, N.Y) 312, 1183–1187.

48. Roure, A., Lemaire, P., and Darras, S. (2014). An Otx/Nodal Regulatory Signature for Posterior Neural Development in Ascidians. PLoS Genetics 10, e1004548. 10.1371/journal.pgen.1004548.

49. Joyce Tang, W., Chen, J.S., and Zeller, R.W. (2013). Transcriptional regulation of the peripheral nervous system in Ciona intestinalis. Dev. Biol. 378, 183–193. 10.1016/j.ydbio.2013.03.016.

50. Waki, K., Imai, K.S., and Satou, Y. (2015). Genetic pathways for differentiation of the peripheral nervous system in ascidians. Nature Communications 6, 8719. 10.1038/ncomms9719.

51. Delsuc, F., Philippe, H., Tsagkogeorga, G., Simion, P., Tilak, M.-K., Turon, X., López-Legentil, S., Piette, J., Lemaire, P., and Douzery, E.J.P. (2018). A phylogenomic framework and timescale for comparative studies of tunicates. BMC Biology 16, 39. 10.1186/s12915-018-0499-2.

52. Kocot, K.M., Tassia, M.G., Halanych, K.M., and Swalla, B.J. (2018). Phylogenomics offers resolution of major tunicate relationships. Molecular Phylogenetics and Evolution 121, 166–173. 10.1016/j.ympev.2018.01.005.

53. Lu, S., Wang, J., Chitsaz, F., Derbyshire, M.K., Geer, R.C., Gonzales, N.R., Gwadz, M., Hurwitz, D.I., Marchler, G.H., Song, J.S., et al. (2020). CDD/SPARCLE: the conserved domain database in 2020. Nucleic Acids Research 48, D265–D268. 10.1093/nar/gkz991.

54. Potter, S.C., Luciani, A., Eddy, S.R., Park, Y., Lopez, R., and Finn, R.D. (2018). HMMER web server: 2018 update. Nucleic Acids Research 46, W200–W204. 10.1093/nar/gky448.

55. Dardaillon, J., Dauga, D., Simion, P., Faure, E., Onuma, T.A., DeBiasse, M.B., Louis, A., Nitta, K.R., Naville, M., Besnardeau, L., et al. (2020). ANISEED 2019: 4D exploration of genetic data for an extended range of tunicates. Nucleic Acids Res 48, D668–D675. 10.1093/nar/gkz955.

